# The “Duckweed Dip”: Aquatic *Spirodela polyrhiza* Plants Can Efficiently Uptake Dissolved, DNA-Wrapped Carbon Nanotubes from Their Environment for Transient Gene Expression

**DOI:** 10.1101/2023.08.21.554121

**Authors:** Tasmia Islam, Swapna Kalkar, Rachel Tinker-Kulberg, Tetyana Ignatova, Eric A. Josephs

## Abstract

Duckweeds (*Lemnaceae*) are aquatic non-grass monocots that are the smallest and fastest-growing flowering plants in the world. While having simplified morphologies, relatively small genomes, and many other ideal traits for emerging applications in plant biotechnology, duckweeds have been largely overlooked in this era of synthetic biology. Here, we report that Greater Duckweed (*Spirodela polyrhiza*), when simply incubated in a solution containing plasmid-wrapped carbon nanotubes (DNA-CNTs), can directly up-take the DNA-CNTs from their growth media with high efficiency and that transgenes encoded within the plasmids are expressed by the plants—without the usual need for large doses of nanomaterials or agrobacterium to be directly infiltrated into plant tissue. This process, called the “duckweed dip”, represents a streamlined, ‘hands-off’ tool for transgene delivery to a higher plant that we expect will enhance the throughput of duckweed engineering and help to realize duckweed’s potential as a powerhouse for plant synthetic biology. (148 words)

---

Duckweeds (*Lemnaceae* or *Araceae*; there are taxonomic disagreements) are small, aquatic, non-grass monocots of which there are 37 species in five genera.^1^ They are the smallest flowering plants in the world, with the largest duckweeds (Greater Duckweed, *Spirodela polyrhiza*) consisting mostly of a small leaf-like structure known as a frond measuring <1 cm across (Figure 1A).^2^ *S. polyrhiza* also has several rhizoid structures underneath their fronds, hence its scientific name.^3-4^ In addition to their simplified morphologies, they also have a significantly reduced genome with fewer than 20,000 genes; several draft genomes and transcriptomic sequencing results have been published. ^5-11^ While they are capable of flowering under certain conditions, in general duckweeds survive in a juvenile state, and *S. polyrhiza* primarily propagates vegetatively by continuously budding off clonal “daughter” plantlets from two meristematic pads asexually every 1 – 3 days (Figure 1A).^5, 12^ Their simplicity to culture in large numbers (with exponential population growth of genetically identical plants) and their ability to efficiently extract material like radiolabeled metabolites from their aquatic environments made them ideal as a model higher plant in the decades before *Arabidopsis*,^13^ but duckweeds have largely been overlooked in this emerging era of synthetic biology, where new approaches have the power to truly unleash their immense biotechnological potential.

**Figure 1.**
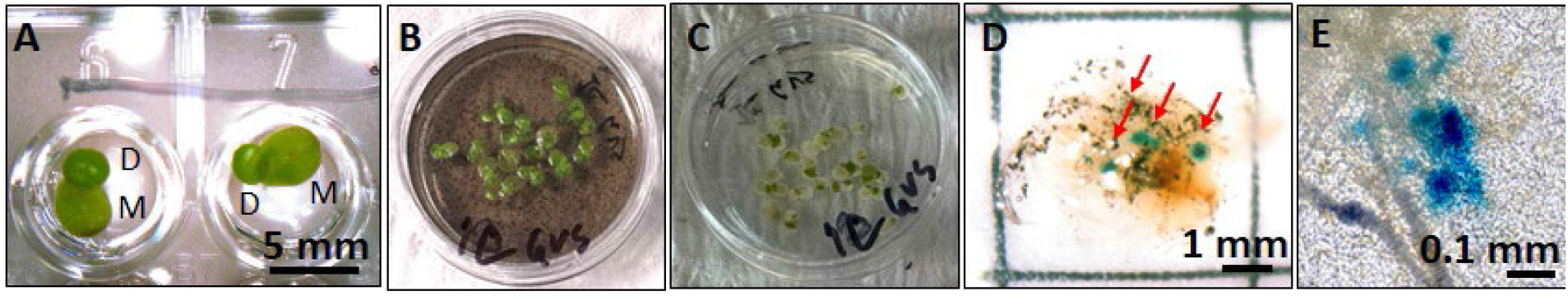
The “duckweed dip”. A) Spirodela polyrhiza mother (M) and daughter (D) fronds growing in the wells of a 96-well plate. B) S. polyrhiza plants are incubated in a 30 mm petri dish containing a solution with carbon nanotubes (DNA-CNTs) wrapped with a plasmid containing the gene for reporter protein β-glucuronidase (GUS) under two 35S promoters. C) After a rinsing / solution exchange to growth media. D) 32 hours after initial incubation in DNA-CNT solution and GUS staining, blue regions on the fronds indicate that S. polyrhiza are positive for in planta GUS expression and activity. E) Closeup of a positive GUS stains in the S. polyrhiza fronds. (all images were brightened 20% for clarity.)

With regards to their biotechnological potential, engineered *Spirodela* species can exhibit very high levels of transgene protein expression: in one report stable, transgenic, GFP yield was >25% of total soluble protein,^14-15^ among the highest in all plant expression systems. Unlike many terrestrial plants, duckweeds can also be sterilized and grown aseptically,^2^ a useful attribute for biopharmaceutical production, and engineered duckweeds have been used to generate monoclonal antibodies and pharmaceutical proteins such as anticoagulants and interferons.^15^ Duckweeds are also edible,^16^ with excellent nutritional properties and biomass accumulation rates comparable to the fastest-growing terrestrial crop plants; engineered duckweeds expressing viral antigens have been used for oral vaccination of animals when given as feed.^17-19^ A particularly powerful recent example of the potential of duckweed synthetic biology was the recent engineering of duckweed *Lemna japonica* for biofuel production:^20^ after inserting a synthetic gene cassette for estradiol-inducible cyan fluorescent protein-*Arabidopsis* WRINKLED1 fusion protein, strong constitutive expression of a mouse diacylglycerol:acyl-CoA acyltransferase 2, and a variant of sesame oleosin, the duckweed exhibited an 108-fold increase in triacylglycerol accumulation that, if grown at scale (on wastewater tracks, for example), would produce 7 times as much oil/biofuel as soy per acre and about as much as oil palm. Applications such as these motivate our work to identify simple, high-efficiency methods to manipulate duckweed biology.

The ability to rapidly screen synthetic gene constructs and cassettes for activity in plants can significantly improve the throughput of phenotype-genotype discovery in plant biology and help to drive success in applications of plant synthetic biology. This can be performed in plants using transient expression systems *via* agrobacterium-mediated infiltration of whole plants, including for *S. polyrhiza*.^21^ However, agrobacterium-mediated approaches tend to have low success rates, can exhibit plant toxicity, and require delicate handling of the duckweeds, including removal of the fragile plantlets from water onto solid media and back again.^21-22^ In terrestrial plants, an alternative strategy for gene delivery that uses DNA-wrapped carbon nanotubes (CNTs) can also be an effective method to deliver plasmid DNA for transient expression.^23-24^ In terrestrial plants, these methods typically require manual infiltration of the nanotubes into plant tissue *via* syringe, which would be difficult in the case of duckweed given their small size and fragility. We hypothesized that we could develop a system whereby simply incubating duckweeds with the DNA-wrapped single-wall CNTs (DNA-CNTs) would result in successful gene delivery because they are so good at extracting material from their aquatic environments. Such a method would allow for stream-lined, high-throughput transient gene expression in duckweeds with minimal “hands-on” time.

In this technical report, we report that, after making some modifications to the reported methods to prepare DNA-loaded SWCNTs that were infiltrated into terrestrial plant tissue, *S. polyrhiza* could survive and were healthy after culture in media containing DNA-CNTs for prescribed times from 30 minutes to at least 48 hours, followed by media exchange / rinsing with water and culture in 0.5 x Schenk & Hildebrandt (SH) media (Figures 1B-1C). We report that after incubation with single-walled CNTs loaded with a plasmid containing the gene for reporter protein β-glucuronidase (GUS) under two 35S promoters,^25^ GUS activity was detected on the fronds of >60% of *S. polyrhiza* plants (at least 15 plants across two technical replicates) after 32 hours (Figures 1D-1E). GUS activity was not detected in plants incubated with plasmid DNA alone (or CNTs alone) and required gentle wounding of the plants with a thin gauge (26G) syringe needle prior to incubation. Plants incubated for 32 hours then washed were also imaged after GUS staining using a confocal Raman microscope,^26-30^ and the strong G-band Raman signature of SWCNTs at 1589 cm^-1^ was detected in frond tissue (Figure 2A-C); no Raman signature of SWCNTS was found in the rhizoid structures (Figure 2D). Thus, we conclude that single-walled CNTs facilitate the passive introduction of plasmid DNA into *S. polyrhiza* fronds where transgenes can be transiently expressed. Detailed methods for this “duckweed dip” protocol, where *S. polyrhiza* are simply incubated in solutions of DNA-loaded SWCNTs followed by media exchange, and which we name in analogy to the simple “floral dip” protocol for transgene delivery used in more easily-transformable higher plants like model *Arabidopsis thaliana*.^31^

**Figure 2.**
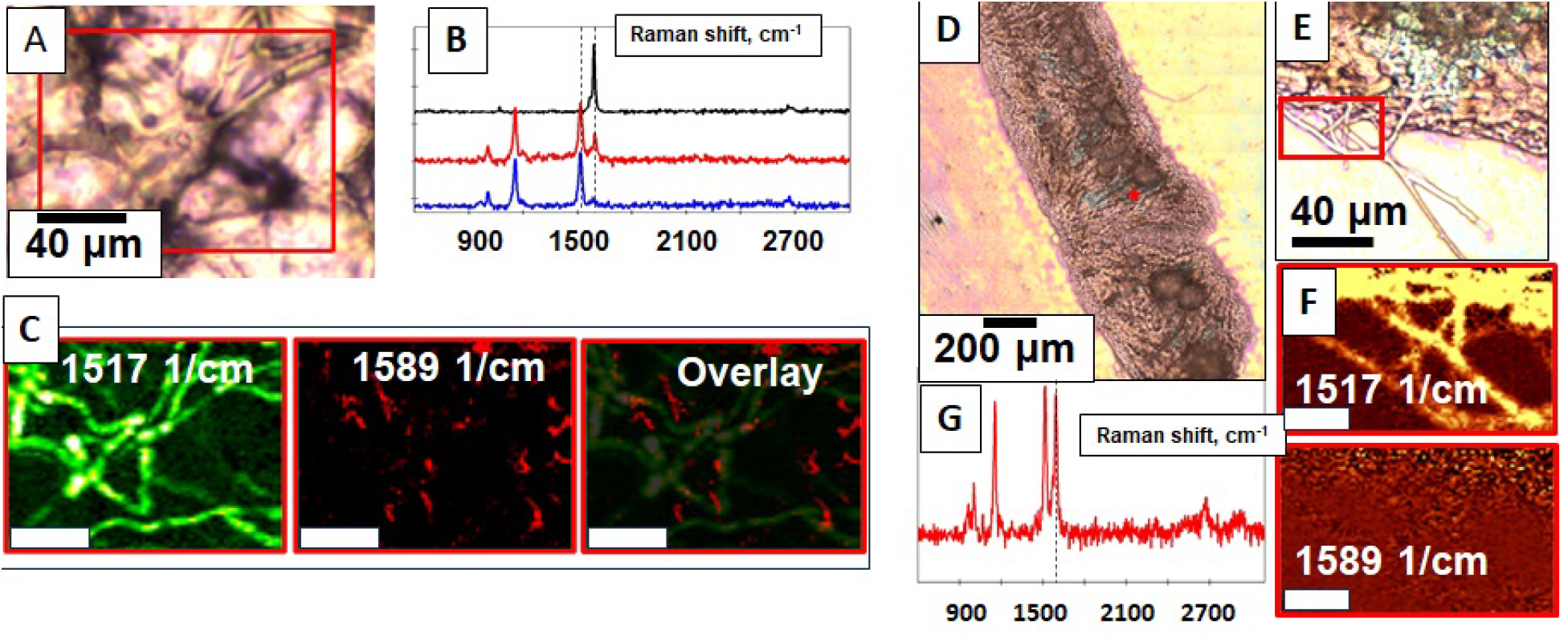
Internalization of functionalized CNTs inside S. polyrhiza. A,D,E) Optical micrographs of duckweed fronds after 32 hr incubation in DNA-CNT solution; B) The Raman spectra showing (black) CNTs alone, (red) duckweed fronds after incubation with DNA-CNTs at the site of blue GUS staining (blue) signal from duckweed tissue away from the sites of blue GUS staining. C) confocal Raman images of (left) duckweed conjugated pigment structures (likely chlorophyll^32^), (middle) CNTs, and (right) overlaid Raman map. G) Raman spectrum taken at the red star location in D) panel shows colocalization of CNT and the region of blue positive GUS staining. F) On the contrast, no CNTs are seen in Raman map of rhizoids (taken in the area highlighted by red square in panel E). Scale in bars C) and F) are 10 μm.

It is notable that *S. polyrhiza*, known for its ability to extract material from their aquatic environments, can not only uptake plasmid-wrapped CNTs but also then express genes encoded on that DNA in the absence of the usual direct delivery of large doses of *Agrobacterium tumefaciens* or DNA-loaded nanomaterials directly into plant tissue. We expect this approach can be applicable to other duckweeds as well. The advantages of our approach reported here are the minimal hands-on time—potentially allowing automation of the transformation process, since only buffer exchanges and gentle wounding are necessary—and the small volumes of materials required for transforming the small plants which can be performed in 96-well plates (Figure 1). Both of those features can greatly facilitate high-throughput plant biology and plant biotechnology optimization. We expect this simple approach to transgene delivery will allow for more efficient duckweed engineering and can serve as a useful tool help to realize duckweed’s strong potential as a powerhouse for plant synthetic biology.

## ASSOCIATED CONTENT

### Supporting Information

Detailed methods and additional plant images are available in the Supporting Information. The Supporting Information is available free of charge on the ACS Publications website.

## Notes

The authors declare no competing financial interests.

## ACKNOWLEDGMENT

We thank Dr. Rammyani Bagchi for expert technical assistance and informative discussions. The work was supported by the National Institutes of Health/National Institute of General Medical Sciences [1R35GM133483] and Department of Defense [Contract #W911QY2220006]. This work was performed in part at the Joint School of Nanoscience and Nanoengineering, a member of the Southeastern Nanotechnology Infrastructure Corridor (SENIC) and National Nanotechnology Coordinated Infrastructure (NNCI), which is supported by the National Science Foundation [ECCS-1542174].

