## Supplementary Information for "The “Duckweed Dip”: Aquatic *Spirodela polyrhiza* Plants Can Efficiently Uptake Dissolved, DNA-Wrapped Carbon Nanotubes from Their Environment for Transient Gene Expression"

**Supporting Information**

**<u>Contents</u>**

**Detailed methods** 2

**Figure S1: Additional images of fronds after the duckweed dip and GUS staining** 4

**Figure S2: Example images of fronds after "control" duckweed dip variations** 5

**Supplementary References** 6

**Detailed methods:**

**Carbon Nanotube Preparation**

The carbon nanotube (CNT) solution was prepared from “as-produced” carboxylic acid functionalized single-wall carbon nanotubes (COOH-SWNTs) (Sigma-Aldrich, cat no. 652490). Dry COOH-SWNTs (15 mg) had been dissolved in 15 mL of nuclease-free water and bath-sonicated for 10 min at 2˚C. Then tip–sonication had been done for 30 min (with a 2-mm probe tip at 10W power) in an ice bath using an ultrasonic homogenizer. After tip sonication the CNT solution had been allowed to rest at room temperature for 10 min then centrifuged at 13,300g for 10 min. The supernatant was withdrawn and again centrifuged for 10 min, then the supernatant withdrawn again. The concentration of COOH-SWNTs was measured *via* absorbance at 632 nm with an extinction coefficient of 0.036 L $mg^{-1}$ $cm^{-1}$. 40 mg of polyethylenimine (PEI, 25,000 MW, branched) was dissolved in 5 mL of 0.1× PBS (pH 7.4–7.6 with 5 M HCl). Then the activated COOH-SWNTs had been added to the PEI solution dropwise. The mixture was then kept for 4 hours at room temperature in an orbital shaker at ~180 r.p.m.

**Plant materials**

*Spirodela polyrhiza 9509* was obtained from Rutgers Duckweed Stock Cooperative (ruduckweed.org). Following collection from the agar plate, the duckweed was surface sterilized in 0.5% sodium hypochlorite and washed with autoclaved water. Duckweed was then cultivated in a practical and inexpensive hydroponic system where plants are grown in a nutrient-rich liquid medium instead of agar. The system consists of two 3-liter stackable bins and a 14-liter plastic reservoir tub. The 3-liter bins were stacked across the larger 14-liter reservoir tub in parallel using the lip of each bin to secure them into place, leaving a 6-inch void space below in the reservoir tub. The 3-liter bins were modified with a nutrient feed pipe through the bottom of each bin that drains into the empty reservoir. A standard submersible Mini-aquarium pump (3W 50 gph) was placed into the bottom reservoir tub and fitted with a Y-shaped tube that allowed water to circulate from the reservoir into the two stacked bins secured on top. UV sterilization of the hydroponic tank to prevent algae growth was provided by placing an aquarium UV lamp (Green Killing Machine, Internal UV) in the bottom reservoir tub as well.

The hydroponic system was filled with commercially sourced nutrient solutions (General Hydroponics FloraSeries Hydroponic Nutrient fertilizer systems with FloraMicro, FloraBloom, and FloraGro) at a 1:1 ratio. Water was circulated (aerated) from the reservoir to the 3-liter bins continuously. Plant density was maintained weekly to prevent overcrowding and renewal of nutrient solution was determined by checking conductivity. Plants in hydroponics tanks were illuminated using a commercially sourced full spectrum LED grow light bulb that was grown 200 mm beneath lamps using a 16 h light/8-h dark cycle. The system was kept under controlled environmental conditions such as nutrients, temperature, light intensity, and photoperiod to optimize growth.

Duckweeds were prepared for experiments by surface sterilization in 2.5% sodium hypochlorite for 1.5 minutes, washed with autoclaved water, then wounded with a fine (26G x 1/2, 0.45 mm x 13 mm) needle on their fronds at ~ 8 spots.

**The “Duckweed Dip”**

Plasmid pSB161 - pL2_pSB90_2x35S::GUS::tMAS, a binary plant vector for transient expression of GUS with introns, was a gift from Erin Cram & Carolyn Lee-Parsons (Addgene plasmid # 123197 ; http://n2t.net/addgene:123197 ; RRID:Addgene_123197). PEI-SWNTs were mixed with this plasmid in a 3:1 PEI-SWNT:DNA mass ratio. 4440 ng of PEI-SWNTs had been diluted in 888 µL of MES delivery buffer and 4090 µLof 0.1x PBS buffer then PEI-SWNTs had been added drop by drop to the 1480 ng (22 µL) of plasmid solution with mixing, then incubated for 30 min at room temperature for the formation of the DNA-PEI-SWNT complex. Then we let the *S. polyrhiza* float atop on that solution with DNA-CNT complexes for 48 hr. After this time, duckweed were briefly rinsed with autoclaved water and maintained in liquid 0.5X SH medium with antibiotic 100 µg/mL Cefatoxime under cool white, fluorescent bulbs in a 16 h light/8-h dark cycle for an additional 3 days.

**GUS reporter assay**

The histological staining of GUS was performed using the protocol described by Yang et al. (2018).[1] Fronds were vacuum infiltrated in staining solution (Sodium phosphate buffer (pH 7.0), 0.5M Ethylene Dia-mine Tetraacetic Acid (EDTA), 50 mM $K3[Fe(CN)_6]$, 50 mM $K4[Fe(CN)_6]$), Triton X (10%), autoclaved water and 0.5 mg/mL 5-bromo-4-chloro-3-indolyl-β-D-glucuronic acid sodium salt (X-Gluc) for 1 h then incubated overnight in the dark at 37 °C. The next day, fronds were washed with 100 mM phosphate buffer (PB) solution and deionized water. Fronds were washed in a solution of 0.24 N HCl and 20% (v/v) methanol for 15 min at 57 °C and another 15 min in a solution of 7% (m/v) NaOH and 10% (v/v) ethanol. Fronds were rinsed with 40% (v/v) ethanol then stored in a solution of 5% (v/v) ethanol and 25% (v/v) glycerin. GUS activity in the fronds of *S. polyrhiza* was imaged using a light microscope. Dark blue spots confirmed the expression of GUS and its activity.

**Raman microscopy**

To confirm the uptake of DNA-CNTs by *S. polyrhiza* plants, we used the WiTec alpha300R Confocal Raman Microscope (Zeiss LD EC Epiplan-Neofluar Dic 50x / 0.55 objective, 600 g/mm grating, 532 nm laser at power not higher than 5W to prevent specimen overheating, integration time 0.1 s) following previously developed protocol [2-4]. The positive *S. polyrhiza* plants were washed and longitudinally sectioned then placed on Si/SiO2 substrate for Raman imaging. DNA-CNTs are localized in sliced specimen by characteristic Raman features on the hyperspectral map, specifically the G (1591 $cm^{-1}$) and G′ (2680 $cm^{-1}$) CNT bands. We also imaged the C = C stretching (1517 $cm^{-1}$) mode which is specific to “skeleton structure” of *S. polyrhiza* plant.

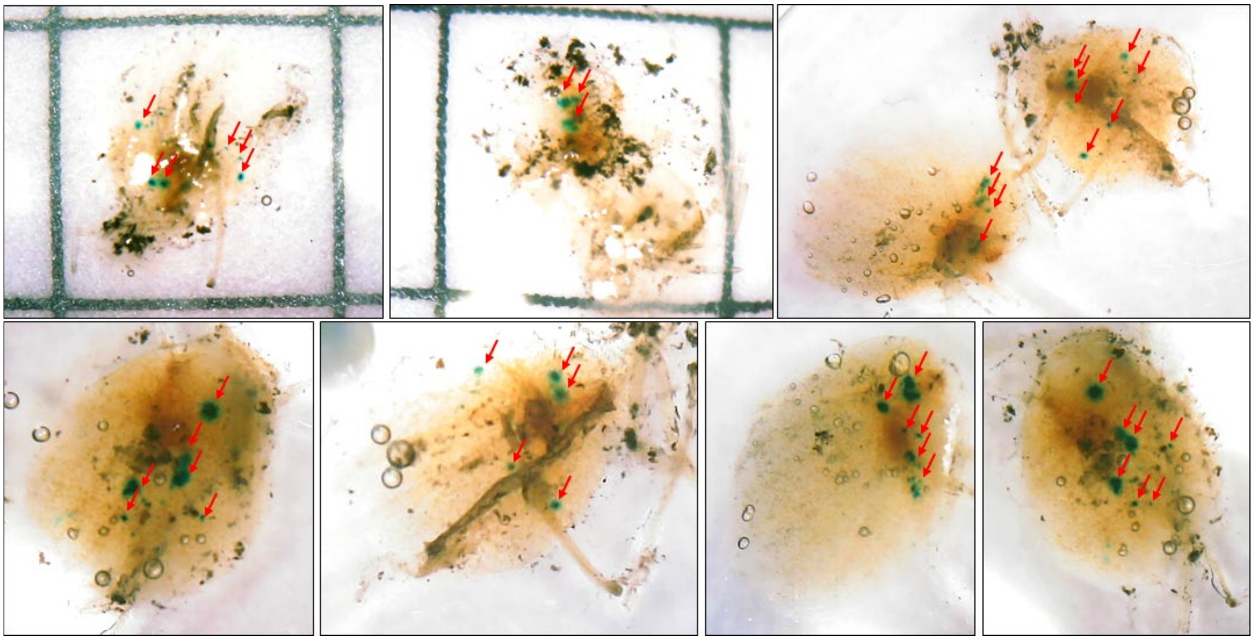

**Figure S1. Additional images of *S. polyrhiza* fronds after the duckweed dip and GUS staining.** Images contain fronds from two technical replicates of the “duckweed dip” protocol. Red arrows highlight blue regions positive for GUS activity. Grid size in top left images is 5 mm. Images were all brightened 20% for clarity.

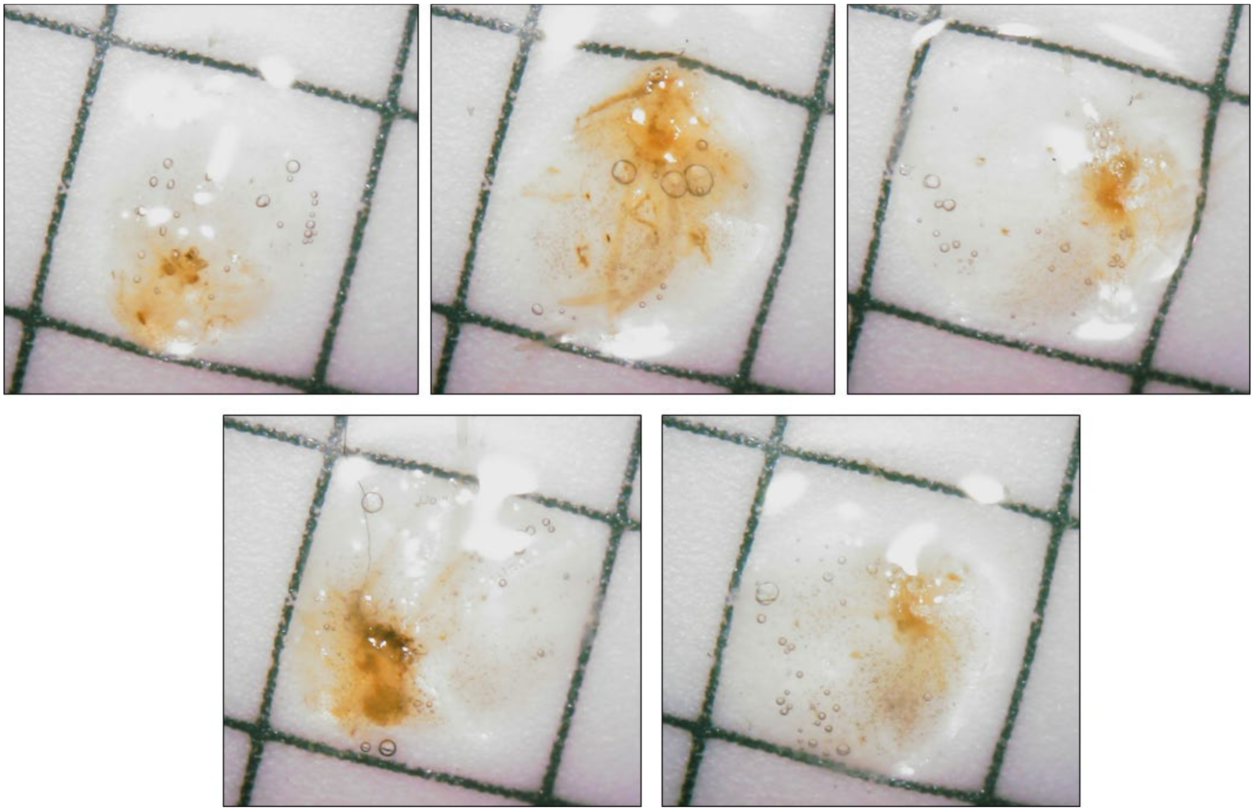

**Figure S2. Example images of fronds after "control" duckweed dip variations.** Duckweeds were "dipped" in a solution containing PEI-SWNTs but no plasmid DNA for 32 hours. After GUS staining, and none exhibit the characteristic blue regions expected for a positive GUS stain as in Figures 1 and S1.
